# Numerical Uncertainty in Analytical Pipelines Lead to Impactful Variability in Brain Networks

**DOI:** 10.1101/2020.10.15.341495

**Authors:** Gregory Kiar, Yohan Chatelain, Oliveira Castro Pablo de, Eric Petit, Ariel Rokem, Gaël Varoquaux, Bratislav Misic, Alan C. Evans, Tristan Glatard

## Abstract

The analysis of brain-imaging data requires complex processing pipelines to support findings on brain function or pathologies. Recent work has shown that variability in analytical decisions, small amounts of noise, or computational environments can lead to substantial differences in the results, endangering the trust in conclusions^1-7^. We explored the instability of results by instrumenting a connectome estimation pipeline with Monte Carlo Arithmetic^8,9^ to introduce random noise throughout. We evaluated the reliability of the connectomes, their features^10,11^, and the impact on analysis^12,13^. The stability of results was found to range from perfectly stable to highly unstable. This paper highlights the potential of leveraging induced variance in estimates of brain connectivity to reduce the bias in networks alongside increasing the robustness of their applications in the classification of individual differences. We demonstrate that stability evaluations are necessary for understanding error inherent to brain imaging experiments, and how numerical analysis can be applied to typical analytical workflows both in brain imaging and other domains of computational science. Overall, while the extreme variability in results due to analytical instabilities could severely hamper our understanding of brain organization, it also leads to an increase in the reliability of datasets.

---

The modelling of brain networks, called connectomics, has shaped our understanding of the structure and function of the brain across a variety of organisms and scales over the last decade^11,14-18^. In humans, these wiring diagrams are obtained *in vivo* through Magnetic Resonance Imaging (MRI), and show promise towards identifying biomarkers of disease. This can not only improve understanding of so-called “connec-topathies”, such as Alzheimer’s Disease and Schizophrenia, but potentially pave the way for therapeutics^19–23^.

However, the analysis of brain imaging data relies on complex computational methods and software. Tools are trusted to perform everything from pre-processing tasks to downstream statistical evaluation. While these tools undoubtedly undergo rigorous evaluation on bespoke datasets, in the absence of ground-truth this is often evaluated through measures of reliability^24–27^, proxy outcome statistics, or agreement with existing theory. Importantly, this means that tools are not necessarily of known or consistent quality, and it is not uncommon that equivalent experiments may lead to diverging conclusions^1,5-7^. While many scientific disciplines suffer from a lack of reproducibility^28^, this was recently explored in brain imaging by a 70 team consortium which performed equivalent analyses and found widely inconsistent results^1^, and it is likely that software instabilities played a role.

The present study approached evaluating reproducibility from a computational perspective in which a series of brain imaging studies were numerically perturbed in such a way that the plausibility of results was not affected, and the implications of the observed instabilities on downstream analyses were quantified. We accomplished this through the use of Monte Carlo Arithmetic (MCA)^8^, a technique which enables characterization of the sensitivity of a system to small numerical perturbations. This is importantly distinct from data perturbation experiments where the underlying datasets are manipulated or pathologies may be simulated, and allows for the evaluation of experimental uncertainty in real-world settings. We explored the impact of numerical perturbations through the direct comparision of structural connectomes, the consistency of their features, and their eventual application in a neuroscience study. We also characterized the consequences of instability in these pipelines on the reliability of derived datasets, and discuss how the induced variability may be harnessed to increase the discriminability of datasets, in an approach akin to ensemble learning. Finally, we make recommendations for the roles perturbation analyses may play in brain imaging research and beyond.

## Graphs Vary Widely With Perturbations

Prior to exploring the analytic impact of instabilities, a direct understanding of the induced variability was required. A subset of the Nathan Kline Institute Rockland Sample (NKIRS) dataset^29^ was randomly selected to contain 25 individuals with two sessions of imaging data, each of which was sub-sampled into two components, resulting in four samples per individual and 100 samples total (25 × 2 × 2 samples). Structural connectomes were generated with canonical deterministic and probabilistic pipelines^30,31^ which were instrumented with MCA, replicating computational noise either sparsely or densely throughout the pipelines^4,9^. In the sparse case, a small subset of the libraries were instrumented with MCA, allowing for the evaluation of the cascading effects of numerical instabilities that may arise. In the dense case, operations are more uniformly perturbed and thus the law of large numbers suggests that perturbations will quickly offset one-another and only dramatic local instabilities will have propagating effects. Importantly, the perturbations resulting from the sparse setting represent a strict subset of the possible outcomes of the dense implementation. The random perturbations are statistically independent from one another across both settings and simulations. Instrumenting pipelines with MCA increases their computation time, in this case by multiplication factors of 1.2× and 7× for the sparse and dense settings, respectively^4^. The results obtained were compared to unperturbed (e.g. reference) connectomes in both cases. The connectomes were sampled 20 times per sample and once without perturbations, resulting in a total of 8, 400 connectomes. Two versions of the unperturbed connectomes were generated and compared such that the absence of variability aside from that induced via MCA could be confirmed.

The stability of connectomes was evaluated through the normalized percent deviation from reference^4^ and the number of significant digits (Figure 1). The comparisons were grouped according to differences across simulations, subsampling of data, sessions of acquisition, or subjects, and accordingly sorted from most to least similar. While the similarity of connectomes decreases as the collections become more distinct, connectomes generated with sparse perturbations show considerable variability, often reaching deviations equal to or greater than those observed across individuals or sessions (Figure 1A; right). Interpretting these results with respect to the distinct MCA environments used suggests that the tested pipelines may not suffer from single dominant sources of instability, but that nevertheless there exist minor local instabilities which may the propagate throughout the pipeline. Furthermore, this finding suggests that instabilities inherent to these pipelines may mask session or individual differences, limiting the trustworthiness of derived connectomes. While both pipelines show similar performance, the probabilistic pipeline was more stable in the face of dense perturbations whereas the deterministic was more stable to sparse perturbations (*p <* 0.0001 for all; exploratory). As an alternative to the normalized percent deviation, the stability of correlations between networks can be found in Supplemental Section S1.

**Figure 1.**
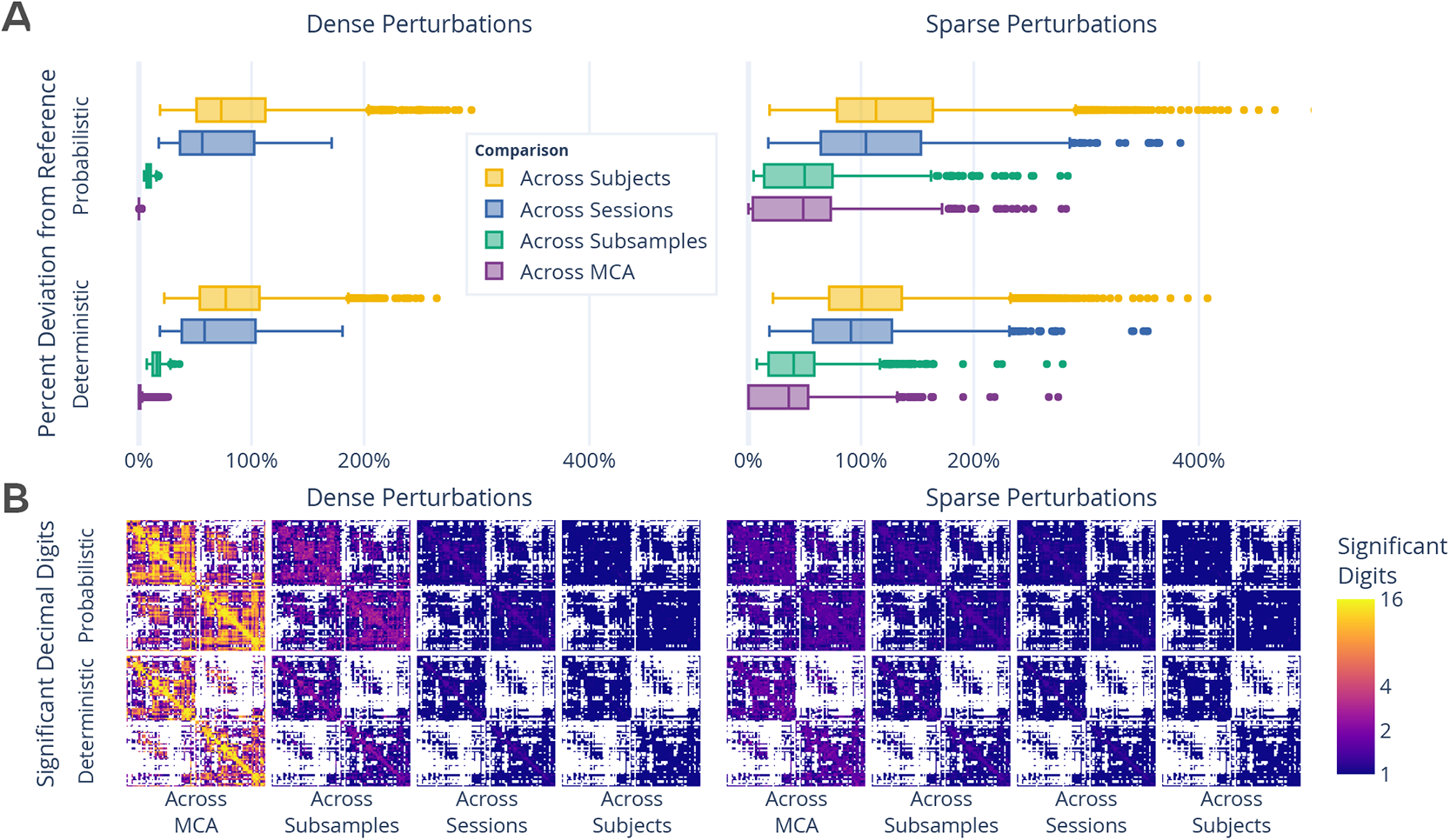
Exploration of perturbation-induced deviations from reference connectomes. (**A**) The absolute deviations between connectomes, in the form of normalized percent deviation from reference. The difference in MCA-perturbed connectomes is shown as the across MCA series, and is presented relative to the variability observed across subsamples, sessions, and subjects. (**B**) The number of significant decimal digits in each set of connectomes as obtained by evaluating the complete distribution of networks. In the case of 16, values can be fully relied upon, whereas in the case of 1 only the first digit of a value can be trusted. Dense and sparse perturbations are shown on the left and right, respectively.

The number of significant digits per edge across connectomes (Figure 1B) similarly decreases alongside the decreasing similarity between comparison groups. While the cross-MCA comparison of connectomes generated with dense perturbations show nearly perfect precision for many edges (approaching the maximum of 15.7 digits for 64-bit data), this evaluation uniquely shows considerable drop off in performance when comparing networks across subsamplings (average of < 4 digits). In addition, sparsely perturbed connectomes show no more than an average of 3 significant digits across all comparison groups, demonstrating a significant limitation in the reliability of independent edge weights. The number of significant digits across individuals did not exceed a single digit per edge in any case, indicating that only the order of magnitude of edges in naively computed groupwise average connectomes can be trusted. The combination of these results with those presented in Figure 1A suggests that while specific edge weights are largely affected by instabilities, macro-scale network structure is stable.

## Sparse Perturbations Reduce Off-Target Signal

We assessed the reproducibility of the dataset through mimicking and extending a typical test-retest experiment^26^ in which the similarity of samples across sessions were compared to distinct samples in the dataset (Table 1, with additional experiments and explanation of the measure and its scaling in Supplemental Section S2). The ability to discriminate connectomes across subjects (Hypothesis 1) is an essential prerequisite for the application of brain imaging towards identifying individual differences^18^. In testing hypothesis 1, we observe that the dataset is discriminable with a scaled score of 0.82 (*p <* 0.001; optimal score: 1.0; chance: 0.04) for both pipelines in the absence of MCA. We can see that inducing instabilities through MCA preserves the discriminability in the dense perturbtion setting, and and discriminability decreased slightly but remained above the unscaled reference value of 0.65 in the sparse case. This lack of significant decrease in discriminability across MCA perturbations suggests its utility for capturing variance within datasets without compromising the robustness and reliability of the dataset as a whole, and possibly suggests this technique as a cost effective and context-agnostic method for dataset augmentation.

**Table 1.** The impact of instabilities as evaluated through the discriminability of the dataset based on individual (or subject) differences, session, and subsample. The performance is reported as mean discriminability. While a perfectly discriminable dataset would be represented by a score of 1.0, the chance performance, indicating minimal discriminability, is 1/the number of classes. *H*_3_ could not be tested using the reference executions due to too few possible comparisons. The alternative hypothesis, indicating significant discrimination, was accepted for all experiments, with *p <* 0.005.

| Comparison | Chance | Target | Unscaled Ref. |  | Scaled Ref. |  | Dense MCA |  | Sparse MCA |  |
| --- | --- | --- | --- | --- | --- | --- | --- | --- | --- | --- |
|  |  |  | Det. | Prob. | Det. | Prob. | Det. | Prob. | Det. | Prob. |
| $H_1$: Across Subjects | 0.04 | 1.0 | 0.64 | 0.65 | 0.82 | 0.82 | 0.82 | 0.82 | 0.77 | 0.75 |
| $H_2$: Across Sessions | 0.5 | 0.5 | 1.00 | 1.00 | 1.00 | 1.00 | 1.00 | 1.00 | 0.88 | 0.85 |
| $H_3$: Across Subsamples | 0.5 | 0.5 | | | | | 0.99 | 1.00 | 0.71 | 0.61 |

While the discriminability of individuals is essential for the identification of individual brain networks, it is similarly reliant on network similarity – or lack of discriminability – across equivalent acquisitions (Hypothesis 2). In this case, connectomes were grouped based upon session, rather than subject, and the ability to distinguish one session from another based on subsamples was computed within-individual and aggregated. Both the unperturbed and dense perturbation settings perfectly preserved differences between sessions with a score of 1.0 (*p <* 0.005; optimal score: 0.5; chance: 0.5), indicating a dominant session-dependent signal for all individuals despite no intended biological differences. However, while still significant relative to chance (score: 0.85 and 0.88; *p <* 0.005 for both), sparse perturbations lead to significantly lower discriminability of the dataset (*p <* 0.005 for all). This reduction of the difference between sessions suggests that the added variance due to perturbations reduces the relative impact of non-biological acquisition-dependent bias inherent in the networks.

Though the previous sets of experiments inextricably evaluate the interaction between data acquisition and tool, the use of subsampling allowed for characterizing the discriminability of networks sampled from within a single acquisition (Hypothesis 3). While this experiment could not be evaluated using reference executions, the networks generated with dense perturbations showed near perfect discrimination between subsamples, with scores of 0.99 and 1.0 (*p <* 0.005; optimal: 0.5; chance: 0.5). Given that there was no variability in data acquisition, due to undesired effects such as participant motion, or preprocessing, the ability to discriminate between equivalent subsamples in this experiment may only be due to instability or bias inherent to the pipelines. The high variability introduced through sparse perturbations considerably lowered the discriminability towards chance (score: 0.71 and 0.61; *p <* 0.005 for all), further supporting this as an effective method for obtaining lower-bias estimates of individual connectivity.

Across all cases, the induced perturbations maintained the ability to discriminate networks on the basis of meaningful biological signal alongside a reduction in discriminability due to of off-target signal in the sparse perturbation setting. This result appears strikingly like a manifestation of the well-known bias-variance tradeoff^32^ in machine learning, a concept which observes a decrease in bias as variance is favoured by a model. In particular, this highlights that numerical perturbations can be used to not only evaluate the stability of pipelines, but that the induced variance may be leveraged for the interpretation as a robust distribution of possible results.

## Distributions of Graph Statistics Are Reliable, But Individual Statistics Are Not

Exploring the stability of topological features of connectomes is relevant for typical analyses, as low dimensional features are often more suitable than full connectomes for many analytical methods in practice^11^. A separate subset of the NKIRS dataset was randomly selected to contain a single non-subsampled session for 100 individuals (100 × 1 × 1) using the pipelines and instrumentation methods to generate connectomes as above. Connectomes were generated 20 times each, resulting in a dataset which also contained 8, 400 connectomes with the MCA simulations serving as the only source of repeated measurements.

The stability of several commonly-used multivariate graph features^10^ were explored and are presented in Figure 2. The cumulative density of the features was computed within individuals and the mean cumulative density and associated standard error were computed for across individuals (Figures 2A and 2B). There was no significant difference between the distributions for each feature across the two perturbation settings, suggesting that the topological features summarized by these multivariate features are robust across both perturbation modes.

**Figure 2.**
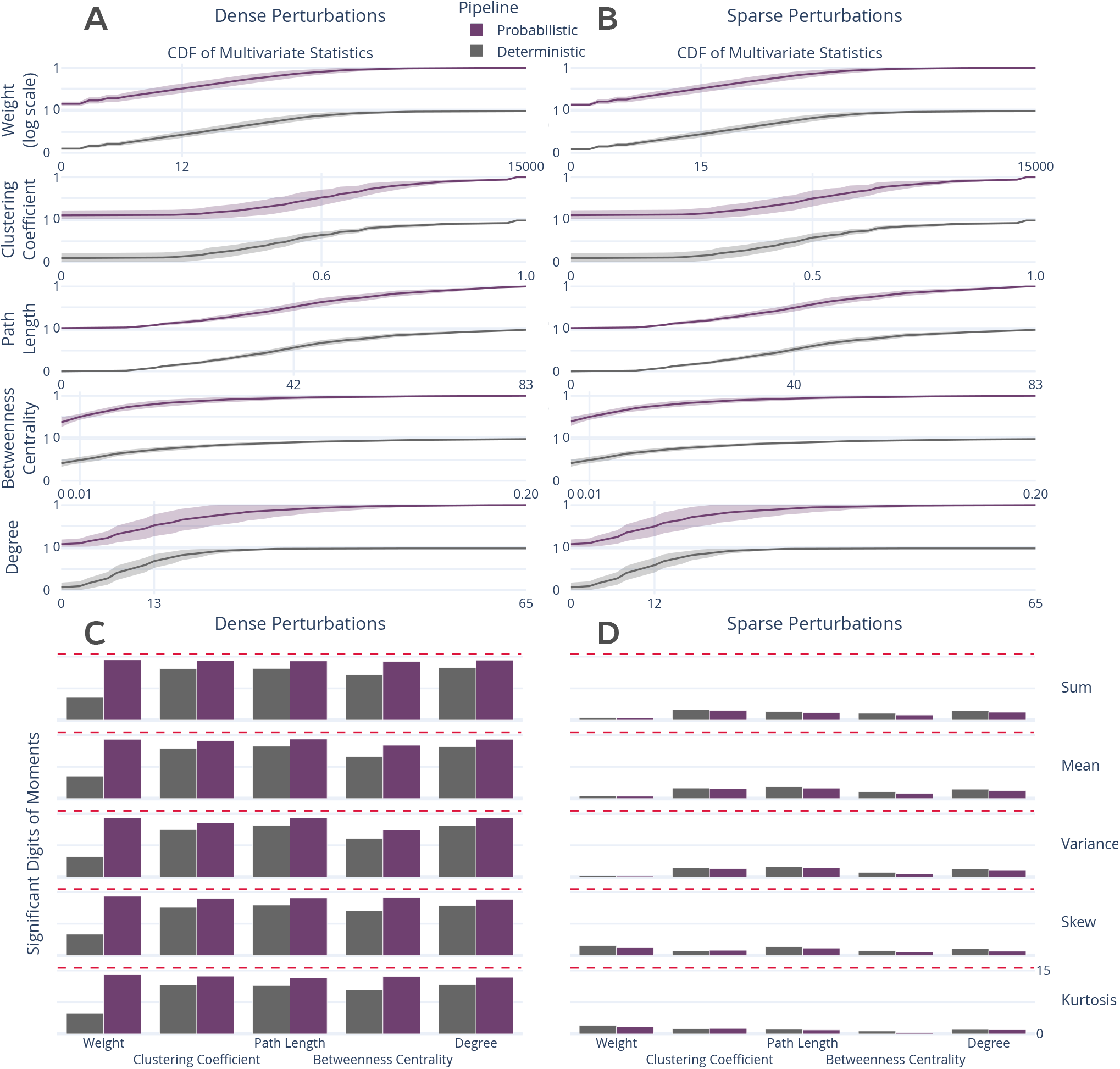
Distribution and stability assessment of multivariate graph statistics. (**A, B**) The cumulative distribution functions of multivariate statistics across all subjects and perturbation settings. There was no significant difference between the distributions in A and B. (**C, D**) The number of significant digits in the first 5 five moments of each statistic across perturbations. The dashed red line refers to the maximum possible number of significant digits.

In addition to the comparison of distributions, the stability of the first 5 moments of these features was evaluated (Figures 2C and 2D). In the face of dense perturbations, the feature-moments were stable with more than 10 significant digits with the exception of edge weight when using the deterministic pipeline, though the probabilistic pipeline was more stable for all comparisons (*p <* 0.0001; exploratory). In stark contrast, sparse perturbations led to highly unstable feature-moments (Figure 2D), such that none contained more than 5 significant digits of information and several contained less than a single significant digit, indicating a complete lack of reliability. This dramatic degradation in stability for individual measures strongly suggests that these features may be unreliable as individual biomarkers when derived from a single pipeline evaluation, though their reliability may be increased when studying their distributions across perturbations. A similar analysis was performed for univariate statistics which obtained similar findings and can be found in Supplemental Section S3.

## Uncertainty in Brain-Phenotype Relationships

While the variability of connectomes and their features was summarized above, networks are commonly used as inputs to machine learning models tasked with learning brain-phenotype relationships^18^. To explore the stability of these analyses, we modelled the relationship between high- or low-Body Mass Index (BMI) groups and brain connectivity using standard dimensionality reduction and classification tools^12^,13, and compared this to reference and random performance (Figure 3).

**Figure 3.**
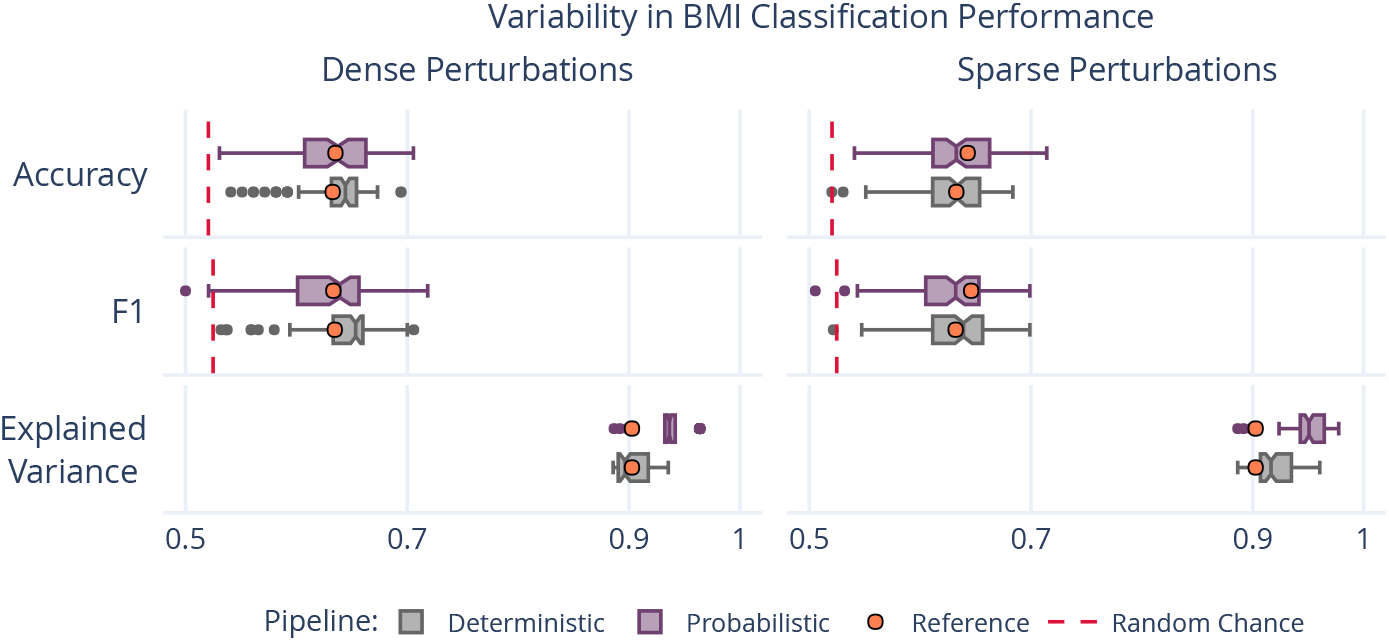
Variability in BMI classification across the sampling of an MCA-perturbed dataset. The dashed red lines indicate random-chance performance, and the orange dots show the performance using the reference executions.

The analysis was perturbed through distinct samplings of the dataset across both pipelines and perturbation methods. The accuracy and F1 score for the perturbed models varied from 0.520 – 0.716 and 0.510 – 0.725, respectively, ranging from at or below random performance to outperforming performance on the reference dataset. This large variability illustrates a previously uncharacterized margin of uncertainty in the modelling of this relationship, and limits confidence in reported accuracy scores on singly processed datasets. The portion of explained variance in these samples ranged from 88.6% -– 97.8%, similar to the reference of 90.3%, suggesting that the range in performance was not due to a gain or loss of meaningful signal, but rather the reduction of bias towards specific outcome. Importantly, this finding does not suggest that modelling brain-phenotype relationships is not possible, but rather it sheds light on impactful uncertainty that must be accounted for in this process, and supports the use of ensemble modeling techniques.

One distinction between the results presented here and the previous is that while networks derived from dense perturbations had been shown to exhibit less dramatic instabilities in general, the results here show similar variability in classification performance across the two methods. This consistency suggests that the desired method of instrumentation may vary across experiments. While sparse perturbations result in considerably more variability in networks directly, the two techniques capture similar variability when relating networks to this phenotypic variable. Given the dramatic reduction in computational overhead, a sparse instrumentation may be preferred when processing datasets for eventual application in modelling brain-phenotype relationships.

## Discussion

The perturbation of structural connectome estimation pipelines with small amounts of noise, on the order of machine error, led to considerable variability in derived brain graphs. Across all analyses the stability of results ranged from nearly perfectly trustworthy (i.e. no variation) to completely unreliable (i.e. containing no trustworthy information). Given that the magnitude of introduced numerical noise is to be expected in computational workflows, this finding has potentially significant implications for inferences in brain imaging as it is currently performed. In particular, this bounds the success of studying individual differences, a central objective in brain imaging^18^, given that the quality of relationships between phenotypic data and brain networks will be limited by the stability of the connectomes themselves. This issue is accentuated through the crucial finding that individually derived network features were unreliable despite there being no significant difference in their aggregated distributions. This finding is not damning for the study of brain networks as a whole, but rather is strong support for the aggregation of networks, either across perturbations for an individual or across groups, over the use of individual estimates.

### Underestimated False Positive Rates

While the instability of brain networks was used here to demonstrate the limitations of modelling brain-phenotype relationships in the context of machine learning, this limitation extends to classical hypothesis testing, as well. Though performing individual comparisons in a hypothesis testing framework will be accompanied by reported false positive rates, the accuracy of these rates is critically dependent upon the reliability of the samples used. In reality, the true false positive rate for a test would be a combination of the reported confidence and the underlying variability in the results, a typically unknown quantity.

When performing these experiments outside of a repeated-measure context, such as that afforded here through MCA, it is impossible to empirically estimate the reliability of samples. This means that the reliability of accepted hypotheses is also unknown, regardless of the reported false positive rate. In fact, it is a virtual certainty that the true false positive rate for a given hypothesis exceeds the reported value simply as a result of numerical instabilities. This uncertainty inherent to derived data is compounded with traditional arguments limiting the trustworthiness of claims^33^, and hampers the ability of researchers to evaluate the quality of results. The accompaniment of brain imaging experiments with direct evaluations of their stability, as was done here, would allow researchers to simultaneously improve the numerical stability of their analyses and accurately gauge confidence in them. The induced variability in derived brain networks may be leveraged to estimate aggregate connectomes with lower bias than any single independent observation, leading to learned relationships that are more generalizable and ultimately more useful.

### Cost-Effective Data Augmentation

The evaluation of reliability in brain imaging has historically relied upon the expensive collection of repeated measurements choreographed by massive cross-institutional consortia^34^,35. The finding that perturbing experiments using MCA both preserved the discriminability of the dataset due to biological signal and decreased the discriminability due to off-target differences across acquisitions and subsamples opens the door for a promising paradigm shift. Given that MCA is data-agnostic, this technique could be used effectively in conjunction with, or in lieu of, realistic noise models to augment existing datasets. While this of course would not replace the need for repeated measurements when exploring the effect of data collection paradigm or study longitudinal progressions of development or disease, it could be used in conjunction with these efforts to decrease the bias of each distinct sample within a dataset. In contexts where repeated measurements are typically collected to increase the fidelity of the dataset, MCA could potentially serve as an alternative solution to capture more biological variability, with the added benefit being the savings of millions of dollars on data collection.

### Shortcomings and Future Questions

Given the complexity of recompiling complex software libraries, pre-processing was not perturbed in these experiments as the instrumentation of the canonical workflow used in diffusion image processing would have added considerable technical complexity and computational overhead to the large set of experiments performed here. Other work has shown that linear registration, a core piece of many elements of pre-processing such as motion correction and alignment, is sensitive to minor perturbations^7^. It is likely that the instabilities across the entire processing workflow would be compounded with one another, resulting in even greater variability. While the analyses performed in this paper evaluated a single dataset and set of pipelines, extending this work to other modalities and analyses, alongside the detection of local sources of instability within pipelines, is of interest for future projects.

This paper does not explore methodological flexibility or compare this to numerical instability. Recently, the nearly boundless space of analysis pipelines and their impact on outcomes in brain imaging has been clearly demonstrated^1^. The approach taken in these studies complement one another and explore instability at the opposite ends of the spectrum, with human variability in the construction of an analysis workflow on one end and the unavoidable error implicit in the digital representation of data on the other. It is of extreme interest to combine these approaches and explore the interaction of these scientific degrees of freedom with effects from software implementations, libraries, and parametric choices.

Finally, it is important to state explicitly that the work presented here does not invalidate analytical pipelines used in brain imaging, but merely sheds light on the fact that many studies are accompanied by an unknown degree of uncertainty due to machine-introduced errors. The presence of unknown error-bars associated with experimental findings limits the impact of results due to increased uncertainty. The desired outcome of this paper is to motivate a shift in scientific computing – both in neuroimaging and more broadly – towards a paradigm that favours the explicit evaluation of the trustworthiness of claims alongside the claims themselves.

## Methods

### Dataset

The Nathan Kline Institute Rockland Sample (NKI-RS)^29^ dataset contains high-fidelity imaging and phenotypic data from over 1, 000 individuals spread across the lifespan. A subset of this dataset was chosen for each experiment to both match sample sizes presented in the original analyses and to minimize the computational burden of performing MCA. The selected subset comprises 100 individuals ranging in age from 6 – 79 with a mean of 36.8 (original: 6 – 81, mean 37.8), 60% female (original: 60%), with 52% having a BMI over 25 (original: 54%).

Each selected individual had at least a single session of both structural T1-weighted (MPRAGE) and diffusion-weighted (DWI) MR imaging data. DWI data was acquired with 137 diffusion directions in a single shell; more information regarding the acquisition of this dataset can be found in the NKI-RS data release^29^.

In addition to the 100 sessions mentioned above, 25 individuals had a second session to be used in a test-retest analysis. Two additional copies of the data for these individuals were generated, including only the odd or even diffusion directions (64 + 9 B0 volumes = 73 in either case) such that the acquired data was evenly represented across both portions, and each subsample represented a realistic complete acquisition. This allowed for an extra level of stability evaluation to be performed between the levels of MCA and session-level variation.

In total, the dataset is composed of 100 subsampled sessions of data originating from 50 acquisitions and 25 individuals for in depth stability analysis, and an additional 100 sessions of full-resolution data from 100 individuals for subsequent analyses.

### Processing

The dataset was preprocessed using a standard FSL^36^ workflow consisting of eddy-current correction and alignment. The MNI152 atlas^37^ was aligned to each session of data via the structural images, and the resulting transformation was applied to the DKT parcellation^38^. Subsampling the diffusion data took place after preprocessing was performed on full-resolution sessions, ensuring that an additional confound was not introduced in this process when comparing between down-sampled sessions. The preprocessing described here was performed once without MCA, and thus is not being evaluated.

Structural connectomes were generated from preprocessed data using two canonical pipelines from Dipy^30^: deterministic and probabilistic. In the deterministic pipeline, a constant solid angle model was used to estimate tensors at each voxel and streamlines were then generated using the EuDX algorithm^31^. In the probabilistic pipeline, a constrained spherical deconvolution model was fit at each voxel and streamlines were generated by iteratively sampling the resulting fiber orientation distributions. In both cases tracking occurred with 8 seeds per 3D voxel and edges were added to the graph based on the location of terminal nodes with weight determined by fiber count.

The random state of both pipelines was fixed for all analyses. Fixing this random state led to entirely deterministic repeated-evaluations of the tools, and allowed for explicit attribution of observed variability to limitations in tool precision as provoked by Monte Carlo simulations, rather than the internal state of the algorithm.

### Perturbations

All connectomes were generated with one reference execution where no perturbation was introduced in the processing. For all other executions, all floating point operations were instrumented with Monte Carlo Arithmetic (MCA)^8^ through Verificarlo^9^. MCA simulates the distribution of errors implicit to all instrumented floating point operations (flop). This rounding is performed on a value *x* at precision *t* by:

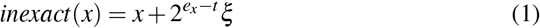

where *e*_*x*_ is the exponent value of *x* and *ξ* is a uniform random variable in the range 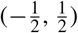. MCA can be introduced in two places for each flop: before or after evaluation. Performing MCA on the inputs of an operation limits its precision, while performing MCA on the output of an operation highlights round-off errors that may be introduced. The former is referred to as Precision Bounding (PB) and the latter is called Random Rounding (RR).

Using MCA, the execution of a pipeline may be performed many times to produce a distribution of results. Studying the distribution of these results can then lead to insights on the stability of the instrumented tools or functions. To this end, a complete software stack was instrumented with MCA and is made available on GitHub at https://github.com/ verificarlo/fuzzy.

The RR variant of MCA was used for all experiments. As was presented in^4^, both the degree of instrumentation (i.e. number of affected libraries) and the perturbation mode have an effect on the distribution of observed results. For this work, the RR-MCA was applied across the bulk of the relevant operations (those occurring in BLAS, LAPACK, Python, Cython, and Numpy) and is referred to as dense perturbation. In this case the bulk of numerical operations were affected by MCA.

Conversely, the case in which RR-MCA was applied across the operations in a small subset of operations (those ocurring in Python and Cython) is here referred to as sparse perturbation. In this case, the inputs to operations within the instrumented libraries were perturbed, resulting in less frequent, data-centric perturbations. Alongside the stated theoretical differences, sparse perturbation is considerably less computationally expensive than dense perturbation.

All perturbations targeted the least-significant-bit for all data (*t* = 24 and *t* = 53 in float32 and float64, respectively^9^). Perturbing the least significant bit importantly serves as a perturbation of machine error, and thus is the appropriate precision to be applied globally in complex pipelines. Simulations were performed 20 times for each pipeline execution for the 100 sample dataset and 10 times for the repeated measures dataset. A detailed motivation for the number of simulations can be found in^39^.

### Evaluation

The magnitude and importance of instabilities in pipelines can be considered at a number of analytical levels, namely: the induced variability of derivatives directly, the resulting downstream impact on summary statistics or features, or the ultimate change in analyses or findings. We explore the nature and severity of instabilities through each of these lenses. Unless otherwise stated, all p-values were computed using Wilcoxon signed-rank tests. To avoid biasing these statistics in this unique repeated-measures context, tests were performed across sets of independent observations and then the results were aggregated in all cases.

### Direct Evaluation of the Graphs

The differences between perturbation-generated graphs was measured directly through both a direct variance quantification and a comparison to other sources of variance such as individual- and session-level differences.

### Quantification of Variability

Graphs, in the form of adjacency matrices, were compared to one another using three metrics: normalized percent deviation, Pearson correlation, and edgewise significant digits. The normalized percent deviation measure, defined in^4^, scales the norm of the difference between a simulated graph and the reference execution (that without intentional perturbation) with respect to the norm of the reference graph, and is defined as^4^:

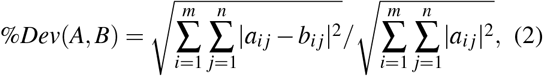

where *A* and *B* each represent a graph, and □_*i j*_ are elements therein corresponding to row and column *i* and *j*, respectively. For these experiments, the *A* graph always refers to the reference, where *B* represents a perturbed value. The purpose of this comparison is to provide insight on the scale of differences in observed graphs relative to the original signal intensity. A Pearson correlation coefficient^40^ was computed in complement to normalized percent deviation to identify the consistency of structure and not just intensity between observed graphs, though the result of this experiment is shown only in Supplemental Section S1.

Finally, the estimated number of significant digits, *s*^*I*^, for each edge in the graph is calculated as:

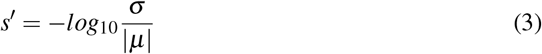

where *µ* and *σ* are the mean and unbiased estimator of standard deviation across graphs, respectively. The upper bound on significant digits is 15.7 for 64-bit floating point data.

The percent deviation, correlation, and number of significant digits were each calculated within a single session of data, thereby removing any subject- and session-effects and providing a direct measure of the tool-introduced variability across perturbations. A distribution was formed by aggregating these individual results.

### Class-based Variability Evaluation

To gain a concrete understanding of the significance of observed variations we explore the separability of our results with respect to understood sources of variability, such as subject-, session-, and pipeline-level effects. This can be probed through Discriminability^26^, a technique similar to ICC^24^ which relies on the mean of a ranked distribution of distances between observations belonging to a defined set of classes. The discriminability statistic is formalized as follows:

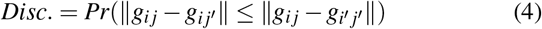

where *g*_*i j*_ is a graph belonging to class *i* that was measured at observation *j*, where *i≠ i*^*I*^ and *j* ≠*j*^*I*^.

Discriminability can then be read as the probability that an observation belonging to a given class will be more similar to other observations within that class than observations of a different class. It is a measure of reproducibility, and is discussed in detail in^26^. This definition allows for the exploration of deviations across arbitrarily defined classes that in practice can be any of those listed above. We combine this statistic with permutation testing to test hypotheses on whether differences between classes are statistically significant in each of these settings. This statistic is similar to *ICC*^24^ in a two-measurement setting, however, given the dependence on a rank distribution from all measurements, discriminability scores do not become meaningless by the addition of more samples which are highly similar to the originals, whereas ICC scores would much more rapidly trend towards 1, making discriminability appropriate in this context. The scaling properties of discriminability are described more fully in Supplemental Section S2.

With this in mind, three hypotheses were defined. For each setting, we state the alternate hypotheses, the variable(s) which were used to determine class membership, and the remaining variables which may be sampled when obtaining multiple observations. Each hypothesis was tested independently for each pipeline and perturbation mode.

#### *H*_*A*1_: Individuals are distinct from one another

Class definition: *Subject ID*

Comparisons: ***Session (1 subsample)***, *Subsample (1 session), MCA (1 subsample, 1 session)*

#### *H*_*A*2_: Sessions within an individual are distinct

Class definition: *Session ID* | *Subject ID*

Comparisons: ***Subsample***, *MCA (1 subsample)*

#### *H*_*A*3_: Subsamples are distinct

Class definition: *Subsample* | *Subject ID, Session ID*

Comparisons: ***MCA***

As a result, we tested 3 hypotheses across 6 MCA experiments and 3 reference experiments on 2 pipelines and 2 perturbation modes, resulting in a total of 30 distinct tests. While results from all tests can be found within Supplemental Section S2, only the bolded comparisons in the list above have been presented in the main body of this article. Correction for repeated testing was performed.

### Evaluating Graph-Theoretical Metrics

While connectomes may be used directly for some analyses, it is common practice to summarize them with structural measures, that can then be used as lower-dimensional proxies of connectivity in so-called graph-theoretical studies^11^. We explored the stability of several commonly-used univariate (graphwise) and multivariate (nodewise or edgewise) features. The features computed and subsequent methods for comparison in this section were selected to closely match those computed in^10^.

### Univariate Differences

For each univariate statistic (edge count, mean clustering coefficient, global efficiency, modularity of the largest connected component, assortativity, and mean path length) a distribution of values across all perturbations within subjects was observed. A Z-score was computed for each sample with respect to the distribution of feature values within an individual, and the proportion of “classically significant” Z-scores, i.e. corresponding to *p <* 0.05, was reported and aggregated across all subjects. There was no correction for multiple comparisons in these statistics, as they were not used to interpret a hypothesis but demonstrate the false-positive rate due to perturbations. The number of significant digits contained within an estimate derived from a single subject were calculated and aggregated. The results of this analysis can be found in Supplemental Section S3.

### Multivariate Differences

In the case of both nodewise (degree distribution, clustering coefficient, betweenness centrality) and edgewise (weight distribution, connection length) features, the cumulative density functions of their distributions were evaluated over a fixed range and subsequently aggregated across individuals. The number of significant digits for each moment of these distributions (sum, mean, variance, skew, and kurtosis) were calculated across observations within a sample and aggregated.

### Evaluating A Brain-Phenotype Analysis

Though each of the above approaches explores the instability of derived connectomes and their features, many modern studies employ modeling or machine-learning approaches, for instance to learn brain-phenotype relationships or identify differences across groups. We carried out one such study and explored the instability of its results with respect to the upstream variability of connectomes characterized in the previous sections. We performed the modeling task with a single sampled connectome per individual and repeated this sampling and modelling 20 times. We report the model performance for each sampling of the dataset and summarize its variance.

### BMI Classification

Structural changes have been linked to obesity in adolescents and adults^41^. We classified normal-weight and overweight individuals from their structural networks (using for overweight a cutoff of BMI > 25^13^). We reduced the dimensionality of the connectomes through principal component analysis (PCA), and provided the first N-components to a logistic regression classifier for predicting BMI class membership, similar to methods shown in^12^,13. The number of components was selected as the minimum set which explained > 90% of the variance when averaged across the training set for each fold within the cross validation of the original graphs; this resulted in a feature of 20 components. We trained the model using *k*-fold cross validation, with *k* = 2, 5, 10, and *N* (equivalent to leave-one-out; LOO).

### Data & Code Provenance

The unprocessed dataset is available through The Consortium of Reliability and Reproducibility (http://fcon_1000. projects.nitrc.org/indi/enhanced/), including both the imaging data as well as phenotypic data which may be obtained upon submission and compliance with a Data Usage Agreement. The connectomes generated through simulations have been bundled and stored permanently (https://doi.org/10.5281/zenodo.4041549), and are made available through The Canadian Open Neuroscience Platform (https://portal.conp.ca/search, search term “Kiar”).

All software developed for processing or evaluation is publicly available on GitHub at https://github.com/ gkpapers/2020ImpactOfInstability. Experiments were launched using Boutiques^42^ and Clowdr^43^ in Compute Canada’s HPC cluster environment. MCA instrumentation was achieved through Verificarlo^9^ available on Github at https://github.com/verificarlo/verificarlo. A set of MCA instrumented software containers is available on Github at https://github.com/gkiar/fuzzy.

## Author Contributions

GK was responsible for the experimental design, data processing, analysis, interpretation, and the majority of writing. All authors contributed to the revision of the manuscript. YC, POC, and EP were responsible for MCA tool development and software testing. AR, GV, and BM contributed to experimental design and interpretation. TG contributed to experimental design, analysis, and interpretation. TG and ACE were responsible for supervising and supporting all contributions made by GK. The authors declare no competing interests for this work. Correspondence and requests for materials should be addressed to Tristan Glatard at tristan.glatard@ concordia.ca.

## Acknowledgments

This research was financially supported by the Natural Sciences and Engineering Research Council of Canada (NSERC) (award no. CGSD3-519497-2018). This work was also supported in part by funding provided by Brain Canada, in partnership with Health Canada, for the Canadian Open Neuroscience Platform initiative.

## S1. Graph Correlation

The following presents a quantification of deviations of generated connectomes from the reference execution, similar to shown in Figure 1. However, in this case, the “percent deviation” measure was replaced with the Pearson correlation coefficient. The correlations between observed graphs (Figure S1) across each grouping follow the same trend to as percent deviation, as shown in Figure 1. However, notably different from percent deviation, there is no significant difference in the correlations between dense or sparse instrumentations. By this measure, the probabilistic pipeline is more stable in all cross-MCA and cross-directions except for the combination of sparse perturbation and cross-MCA (*p* < 0.0001 for all; exploratory).

The marked lack in drop-off of performance across these settings, inconsistent with the measures show in Figure 1 is likely due to the nature of the measure and the structure of graphs being compared. Given that structural graphs are sparse and contain considerable numbers of zero-weighted edges, the presence or absense of edges dominated the correlation measure where it was less impactful for the others. For this reason and others44, correlation is not a commonly used measure in the context of structural connectivity, and thus this analysis was demoted to the supplement material.

**Figure S1.**
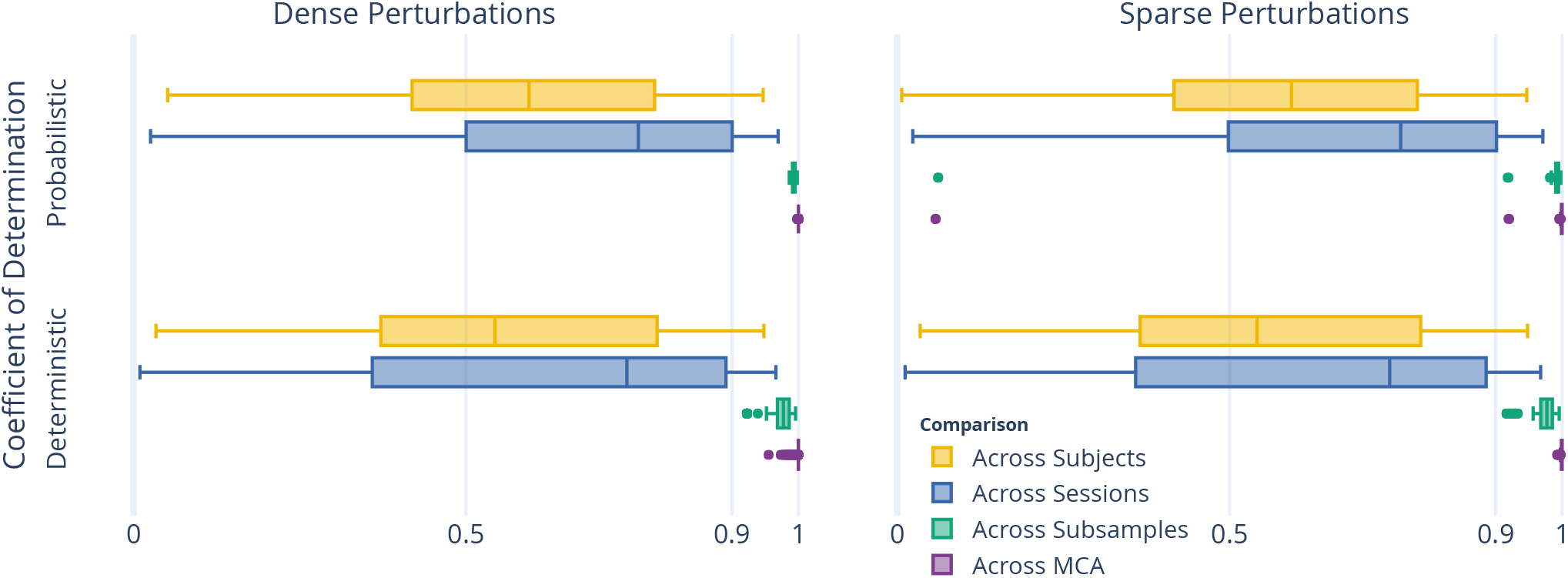
The correlation between perturbed connectomes and their reference.

## S2. Complete Discriminability Analysis

**Table S1.** The complete results from the Discriminability analysis, with results reported as mean *±* standard deviation Discriminability. As was the case in the condensed table, the alternative hypothesis, indicating significant separation across groups, was accepted for all experiments, with *p <* 0.005.

| Exp. | Subj. | Sess. | Samp. | Unscaled Reference |  | Dense Perturbations |  | Sparse Perturbations |  |
| --- | --- | --- | --- | --- | --- | --- | --- | --- | --- |
|  |  |  |  | Det. | Prob. | Det. | Prob. | Det. | Prob. |
| 1.1 | All | All | 1 | $0.64 \pm 0.00$ | $0.65 \pm 0.00$ | $0.82 \pm 0.00$ | $0.82 \pm 0.00$ | $0.77 \pm 0.00$ | $0.75 \pm 0.00$ |
| 1.2 | All | 1 | All | $1.00 \pm 0.00$ | $1.00 \pm 0.00$ | $1.00 \pm 0.00$ | $1.00 \pm 0.00$ | $0.93 \pm 0.02$ | $0.90 \pm 0.02$ |
| 1.3 | All | 1 | 1 | | | $1.00 \pm 0.00$ | $1.00 \pm 0.00$ | $0.94 \pm 0.02$ | $0.90 \pm 0.02$ |
| 2.4 | 1 | All | All | $1.00 \pm 0.00$ | $1.00 \pm 0.00$ | $1.00 \pm 0.00$ | $1.00 \pm 0.00$ | $0.88 \pm 0.12$ | $0.85 \pm 0.12$ |
| 2.5 | 1 | All | 1 | | | $1.00 \pm 0.00$ | $1.00 \pm 0.00$ | $0.89 \pm 0.11$ | $0.84 \pm 0.12$ |
| 3.6 | 1 | 1 | All | | | $0.99 \pm 0.03$ | $1.00 \pm 0.00$ | $0.71 \pm 0.07$ | $0.61 \pm 0.05$ |

The complete discriminability analysis includes comparisons across more axes of variability than the condensed version. The reduction in the main body was such that only axes which would be relevant for a typical analysis were presented. Here, each of Hypothesis 1, testing the difference across subjects, and 2, testing the difference across sessions, were accompanied with additional comparisons to those shown in the main body.

### Subject Variation

Alongside experiment 1.1, that which mimicked a typical test-retest scenario, experiments 1.2 and 1.3 could be considered a test-retest with a handicap, given a single aqcuisition per individual was compared either across subsamples or simulations, respectively. For this reason, it is unsurprising that the dataset achieved considerably higher discriminability scores.

### Session Variation

Similar to subject variation, the session variation was also modelled across either both or a single subsample in experiments 2.4 and 2.5. In both of these cases the performance was similar, and the finding that sparse perturbations reduced the off-target signal was consistent.

### S2.1 Scaling of discriminability with N

When samples were added to the dataset across perturbed executions, the discriminability statistic inflated to a plateau even when no information was added (e.g. the dataset was replicated). This effect is demonstrated for the reference executions and is shown in Figure S2. As we can see, the reference discriminability scores without data duplication (unscaled) were 0.64 and 0.65 for the deterministic and probabilistic pipelines, respectively. After duplicating the dataset 20 times, matching the size of the 20-sample perturbed dataset, we can see that this (scaled) score plateaus at 0.82 for both pipelines. For consistency, in the main body of the text the reference execution performance was communicated as the scaled quantity.

**Figure S2.**
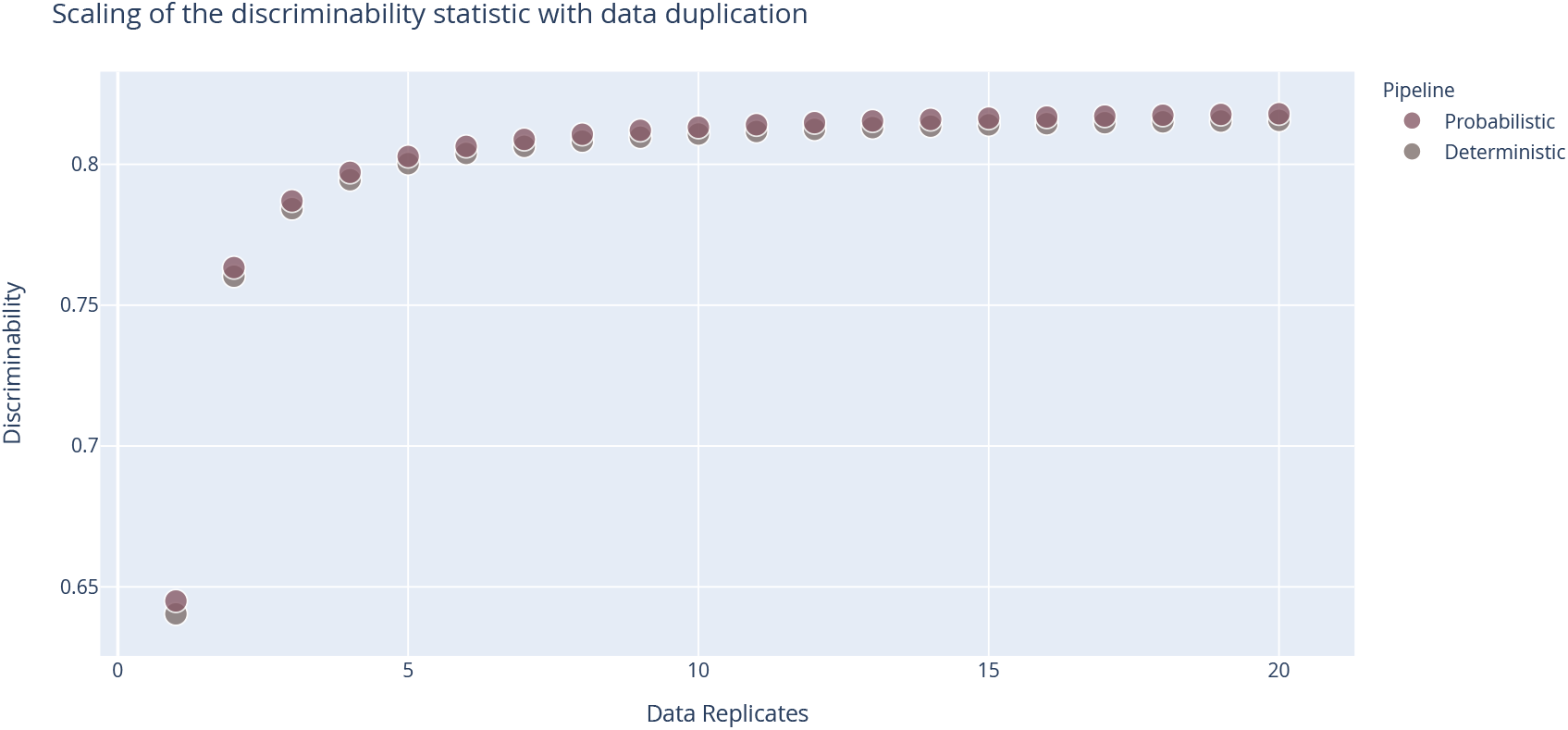
Scaling behaviour of the discriminability statistic with data duplication.

## S3. Univariate Graph Statistics

Figure S3 explores the stability of univariate graph-theoretical metrics computed from the perturbed graphs, including modularity, global efficiency, assortativity, average path length, and edge count. When aggregated across individuals and perturbations, the distributions of these statistics (Figures S3A and S32B) showed no significant differences between perturbation methods for either deterministic or probabilistic pipelines, consistent with the comparison of the cumulative density of the multivariate statistics compared in 2.

However, when quantifying the stability of these measures across connectomes derived from a single session of data, the two perturbation methods show considerable differences. The number of significant digits in univariate statistics for dense perturbation instrumented connectome generation exceeded 11 digits for all measures except modularity, which contained more than 4 significant digits of information (Figure S3C). When detecting false-positives from the distributions of observed statistics for a given session, the rate (using a threshold of *p* = 0.05) was approximately 2% for all statistics with the exception of modularity which again was less stable with an approximately 10% false positive rate. The probabilistic pipeline is significantly more stable than the deterministic pipeline (*p* < 0.0001; exploratory) for all features except modularity. When similarly evaluating these features from connectomes generated in the sparse perturbation setting, no statistic was stable with more than 3 significant digits or a false positive rate lower than nearly 6% (Figure S3D). The deterministic pipeline was more stable than the probabilistic pipeline in this setting (*p* < 0.0001; exploratory).

Two notable differences between the two perturbation methods are, first, the uniformity in the stability of the statistics, and second, the dramatic decline in stability of individual statistics in the sparse perturbation setting despite the consistency in the overall distribution of values. This result is consistent with that obtained from the multivariate exploration performed in the body of this article. It is unclear at present if the discrepancy between the stability of modularity in the pipeline perturbation context versus the other statistics suggests the implementation of this measure is the source of instability or if it is implicit to the measure itself. The dramatic decline in the stability of features derived from sparse perturbed graphs despite no difference in their overall distribution both shows that while individual estimates may be unstable the comparison between aggregates or groups may be considered much more reliable; this finding is consistent with that presented for multivariate statistics.

**Figure S3.**
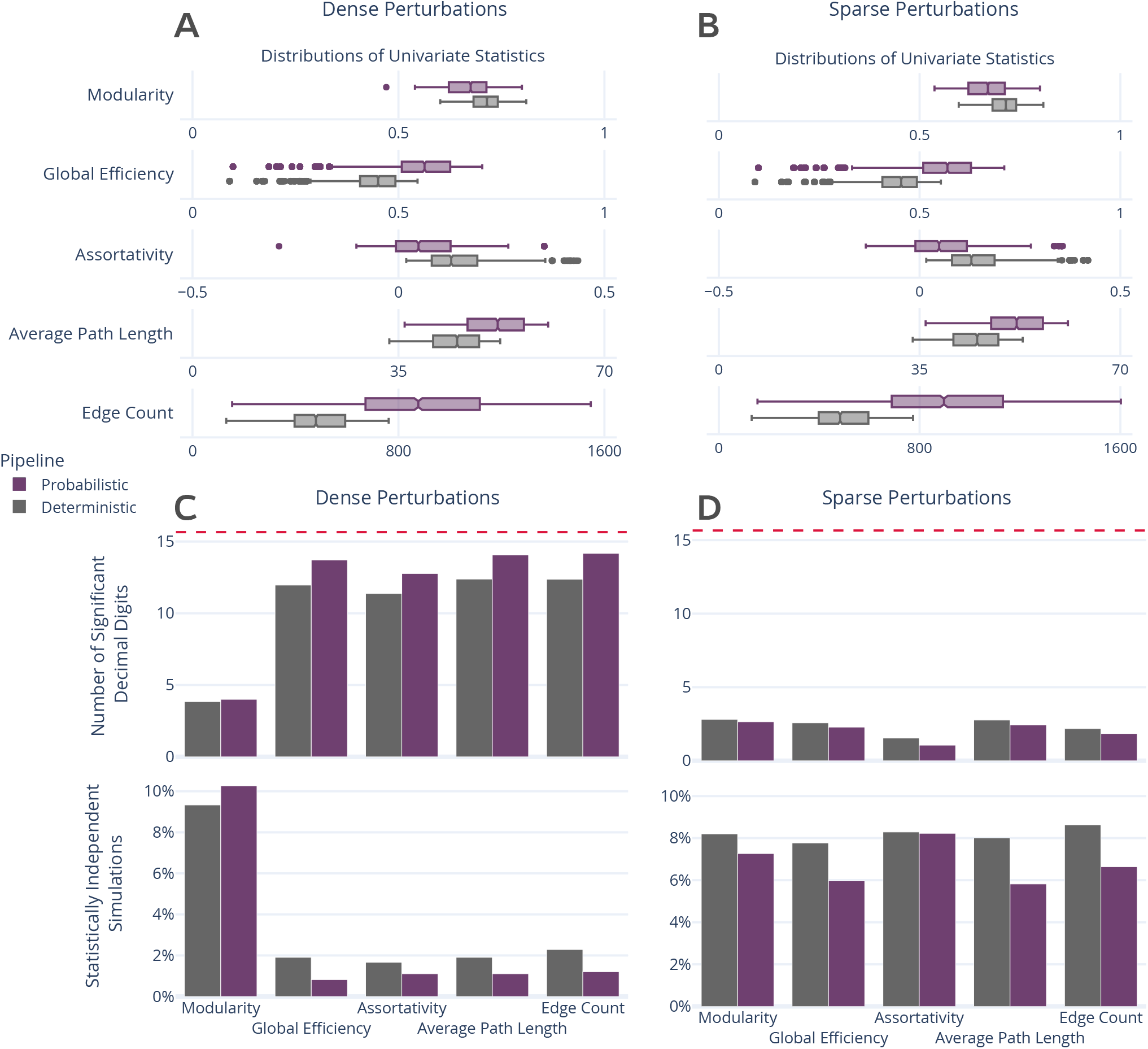
Distribution and stability assessment of univariate graph statistics. (**A, B**) The distributions of each computed univariate statistic across all subjects and perturbations for dense and sparse settings, respectively. There was no significant difference between the distributions in A and B. (**C, D**; top) The number of significant decimal digits in each statistic across perturbations, averaged across individuals. The dashed red line refers to the maximum possible number of significant digits. (**C, D**; bottom) The percentage of connectomes which were deemed significantly different (*p* < 0.05) from the others obtained for an individual.

